# Local adaptation, geographical distance and phylogenetic relatedness: assessing the drivers of siderophore-mediated social interactions in natural bacterial communities

**DOI:** 10.1101/2020.12.16.423016

**Authors:** Elena Butaitė, Jos Kramer, Rolf Kümmerli

**Affiliations:** Department of Plant and Microbial Biology, University of Zurich, Winterthurerstrasse 190, 8057 Zurich, Switzerland; Department of Quantitative Biomedicine, University of Zurich, Winterthurerstrasse 190, 8057 Zurich, Switzerland

**Keywords:** microbe-microbe interactions, siderophores, public goods, genetic isolation-by-distance, Pseudomonas

## Abstract

In heterogenous, spatially structured habitats, individuals within populations can become adapted to the prevailing conditions in their local environment. Such local adaptation has been reported for animals and plants, and for pathogens adapting to hosts. There is increasing interest in applying the concept of local adaptation to microbial populations, especially in the context of microbe-microbe interactions. Here, we tested whether cooperation and cheating on cooperation can spur patterns of local adaptation in soil and pond communities of *Pseudomonas* bacteria, collected across a geographical scale of 0.5 to 50 meters. We focused on the production of pyoverdines, a group of secreted iron-scavenging siderophores that often differ among pseudomonads in their chemical structure and the receptor required for their uptake. A combination of supernatant-feeding and competition assays between isolates from four distance categories revealed tremendous variation in the extent to which pyoverdine non- and low-producers can benefit from pyoverdines secreted by producers. However, this variation was not explained by geographical distance, but primarily depended on the phylogenetic relatedness between interacting isolates. A notable exception occurred in local pond communities, where the effect of phylogenetic relatedness was eroded in supernatant assays, probably due to the horizontal transfer of receptor genes. While the latter result could be a signature of local adaptation, our results overall indicate that common ancestry and not geographical distance is the main predictor of siderophore-mediated social interactions among pseudomonads.

## INTRODUCTION

Local adaptation in spatially structured environments, where dispersal of individuals is limited and environmental patches differ in conditions, is a common phenomenon in animals and plants (Kawecki and Ebert 2004; Leimu and Fischer 2008; Hereford 2009; Blanquart et al. 2013; Savolainen et al. 2013). The concept implies that divergent selection causes local populations to adapt to abiotic and biotic conditions in their respective habitat patch. As a consequence, individuals cope well with prevailing local conditions, but potentially perform poorly (i.e. are mal-adapted) in other patches where conditions differ.

In contrast to animals and plants, the extent to which local adaptation occurs in microbial populations is much less well understood (Giraud et al. 2017; Kraemer and Boynton 2017). The majority of work in this field focused on host-pathogen interactions and examined whether microbial pathogens (i.e. eukaryotic parasites, bacteria, phages) are locally adapted to their animal hosts (Kaltz and Shykoff 1998; Greischar and Koskella 2007). Even fewer studies are available on local adaptation to abiotic conditions (Kraemer and Boynton 2017), with the existing body of work yielding mixed results. For example, Belotte et al. (2003) found significant patterns of local growth adaptation among a collection of *Bacillus* soil isolates across distances of up to 80 m, while others could not confirm such patterns for *Pseudomonas* isolates sampled across a geographical scale of 10-1000 m (Kraemer and Kassen 2015; 2016).

In addition to abiotic conditions, there is an increasing interest in understanding whether bacteria can locally adapt to other members of their community (Kraemer and Boynton 2017). The reason is that bacteria possess numerous traits to socially interact with other microbes (West et al. 2007). Social interactions can include both cooperative interactions, e.g. through the sharing of secreted enzymes or nutrient scavenging molecules (Asfahl and Schuster 2017; Abisado et al. 2018; Kramer et al. 2020b) and competitive interactions, e.g. through the secretion of toxins (Hibbing et al. 2010; Abrudan et al. 2015; Ghoul and Mitri 2016). Microbial toxins such as bacteriocins and antibiotics exert narrow-range activities against closely related strains with considerable niche overlap. Their deployment could hence spur patterns of local adaptation so that bacteria are particularly efficient in combatting their local competitors (Hawlena et al. 2010; Kinkel et al. 2014; Bruce et al. 2017b). However, results on local adaptation with regard to toxin-mediated social interactions are scarce and mixed. While some studies revealed significant geographical distance effects in *Xenorhabdus* and *Streptomyces* populations collected from soil (Hawlena et al. 2010; Kinkel et al. 2014), with inhibition being most pronounced against strains from distant patches, other studies found no or very weak evidence for local adaptation in *Pseudomonas* soil populations (Bruce et al. 2017b; Kraemer et al. 2017). The latter studies further found that strain inhibition via bacteriocins was low overall (< 10%). Bacteriocins and their immunity proteins are often encoded on mobile genetic elements (Nogueira et al. 2009; Silby et al. 2011; Brockhurst et al. 2019), and can thus spread quickly as selfish genetic elements through populations, which might explain both the low frequency of inhibition and the absence of local adaptation.

In our study, we examine whether cooperative interactions and cheating on a cooperative trait can trigger patterns of local adaptation in bacterial communities. We focus on pyoverdine, an iron-chelating siderophore produced by *Pseudomonas* bacteria. Siderophore production can be a cooperative trait because the molecules are secreted in the environment where they scavenge iron from natural sources, making it bioavailable for community members in the vicinity that possess a corresponding receptor for the uptake of the chelated iron (Kramer et al. 2020b). There is increasing evidence that the sharing of siderophores and their exploitation by siderophore non-producers drive community dynamics in marine, fresh-water and soil environments (Cordero et al. 2012; Bruce et al. 2017a; Butaitė et al. 2017; Gu et al. 2020; Kramer et al. 2020b). Given the strong fitness effects siderophores can have, local adaption to siderophore use and exploitation could manifest in two different ways. First, siderophore non-producers could become locally adapted to efficiently exploit siderophores produced by members of their own community. Second, producers could evolve strategies to minimize / avoid being exploited by local non-producers.

At the mechanistic level, such local adaptation could arise due to variation in the chemical structure of siderophores and their corresponding receptors required for uptake (Faraldo-Gómez and Sansom 2003; Hider and Kong 2010; Kümmerli et al. 2014). In this respect, pyoverdine, the main siderophore produced by fluorescent pseudomonads, is an ideal model trait, because many different varieties of this molecule and its receptor exist (Ghysels et al. 2004; Smith et al. 2005; Meyer et al. 2008; Butaitė et al. 2017). While pyoverdines feature a conserved fluorophore, their peptide backbone can vary substantially and is usually strain-specific (Visca et al. 2007; Meyer et al. 2008; Schalk and Guillon 2013). It was suggested that the observed diversity could be the result of antagonistic co-evolution, whereby pyoverdine non-producers that experience a relative fitness advantage by exploiting pyoverdine produced by others, i.e. cheaters, trigger selection for novel, less-exploitable variants of pyoverdines. This process could in turn lead to selection for novel, more efficient cheater types (Smith et al. 2005; Lee et al. 2012; Butaitė et al. 2017). In structured populations with limited dispersal, such antagonistic co-evolution could occur within patches and then give rise to cheaters that are proficient in exploiting local producers, but might be less efficient in exploiting producers from more distant patches with which they have not interacted before.

To explore this scenario, we isolated a total of 315 *Pseudomonas* strains from eight soil and eight pond samples (henceforth called ‘communities’, each comprising 18-20 isolates) across a geographical scale of 50 m for each habitat (Fig. 1A). We first identified pyoverdine non-producers and low-producers (i.e. potential cheaters, henceforth called NLPs) and then tested in a common-garden experiment whether there are differences in the extent to which these NLPs can use the pyoverdine from producers originating either from their own community or different communities at close (0.5 m), intermediate (5 m), and far (50 m) distances. First, we measured the absolute growth benefits NLPs can obtain when being exposed to pyoverdine in supernatants from producers. Second, we measured the relative fitness of NLPs in direct competition with the producers. Moreover, we used sequence data from the *rpoD* gene to test whether there is genetic isolation-by-distance between interacting isolates across the geographical scale sampled, and whether patterns of pyoverdine-mediated interactions correlate with the genetic relatedness between the interacting isolates.

**Figure 1.**
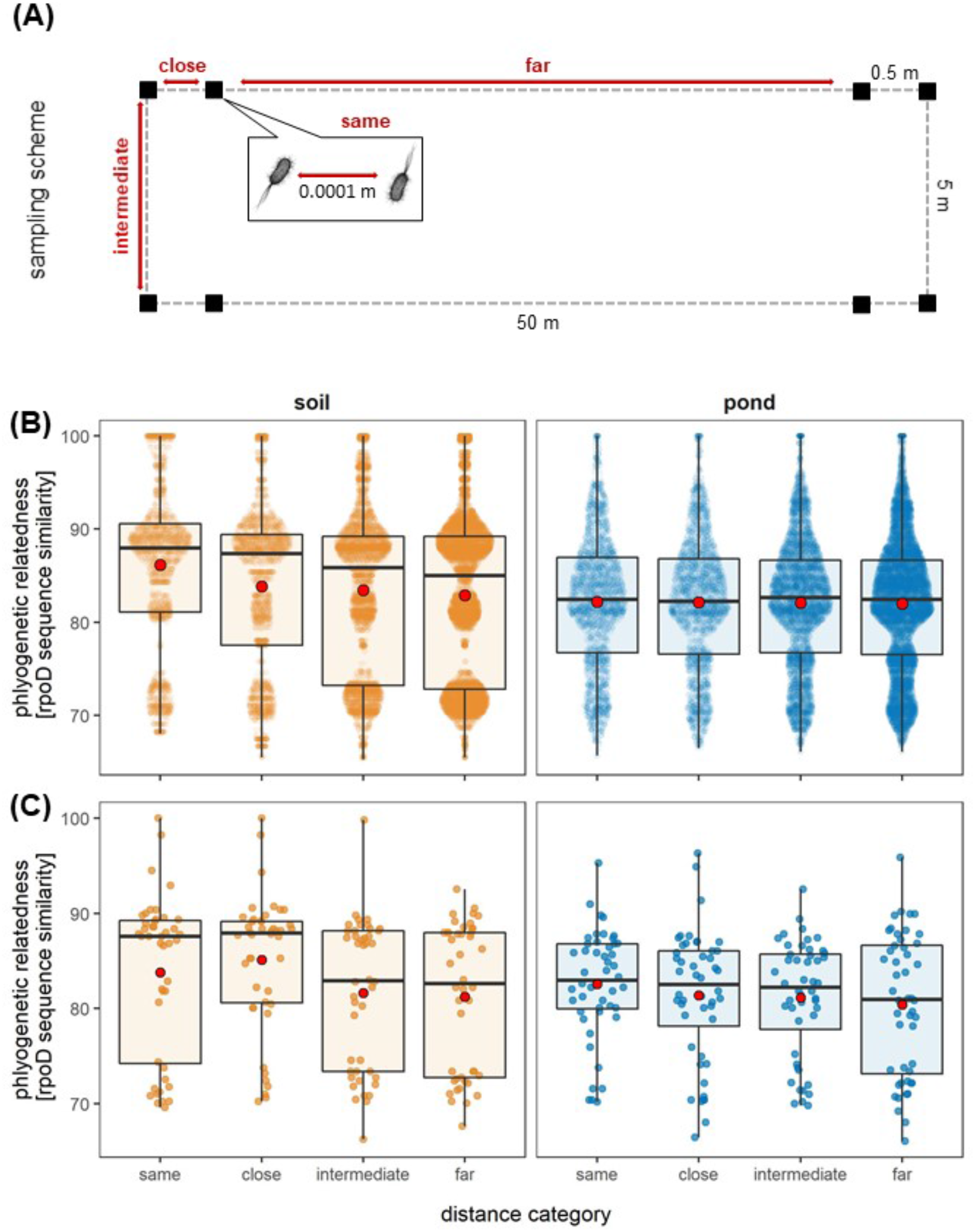
Sampling design and phylogenetic relatedness by distance. (A) Scheme of the sampling design used for both the soil and the pond habitat. Black squares represent the eight independent soil and water communities sampled, and the distances between them. (B) Phylogenetic relatedness between the full collection of soil (yellow) and pond (blue) isolates across the four different distance categories, based on the pairwise comparison of partial *rpoD* sequences. Phylogenetic relatedness significantly declined over distance in the soil but not in the pond. Data were analyzed with a Mantel correlation test for which the distance between isolates from the same community had to be set to a small value larger than zero (i.e. 0.0001 m). (C) Phylogenetic relatedness between the isolates used in the supernatant assay: pyoverdine producers versus pyoverdine NLPs.

## MATERIALS AND METHODS

### Sampling and isolation of pseudomonads

We sampled eight soil cores and eight water samples from a meadow and a pond on the Irchel campus of the University of Zurich (47.40° N, 8.54° E), Switzerland. We followed a rectangular collection scheme (Fig. 1A), such that each soil core and water sample had one close (50 cm apart), two intermediate (5 m) and four distant (50 m) neighboring samples. The isolation and characterization of fluorescent *Pseudomonas* spp. from these samples are described in detail in two previous papers (Butaitė et al. 2017; Butaitė et al. 2018). In brief, bacterial suspensions from soil and pond extracts were plated on Gould’s S1 medium supplemented with 100 μg/ml of the antifungal cycloheximide and 50 μM FeCl3 (to also allow siderophore non-producers to grow). This medium is selective for fluorescent *Pseudomonas* (Gould et al. 1985). Following three days of incubation at room temperature, we randomly picked 20 isolates for each of the eight soil and eight pond samples, 320 isolates in total, re-streaked them on lysogeny broth (LB) agar, and finally picked a single colony for freezer stock preservation at −80°C in a 25%-glycerol-LB mix. Each isolate was provided with an ID code and isolates from the same sample were considered to belong to the same bacterial community.

### Genetic characterization of isolates and phylogenetic analysis

We have previously sequenced the housekeeping gene *rpoD* of all 320 isolates (see Butaitė et al. 2017, for detailed protocols and primers). PCR amplification and sequencing were successful for 315 isolates, while it failed in the remaining five strains, which consequently had to be excluded from further analysis. The *rpoD* gene is commonly used for phylogenetic affiliation of pseudomonads (Mulet et al. 2009; Ghyselinck et al. 2013). We performed in-depth phylogenetic analysis elsewhere (Butaitė et al. 2017; Kramer et al. 2020a), confirming that our 315 isolates belong to the group of fluorescent pseudomonads. For this paper, we estimated the phylogenetic relatedness between interacting strains by carrying out a multiple sequence alignment and obtaining a pairwise identity matrix for 513 bp *rpoD* sequences of 304 strains, using MAFFT (Madeira et al. 2019). Eleven isolates had to be excluded at this stage because their sequence lengths were too short, such that their inclusion would have compromised the resolution in calculating the relatedness. All *rpoD* sequences are deposited at the European Nucleotide Archive (http://www.ebi.ac.uk/ena) under the study accession number PRJEB21289.

### Experimental pairing of isolates

In our previous paper, we screened all isolates for their ability to produce pyoverdine and to grow under iron-limited conditions (Butaitė et al. 2017). Specifically, we made use of the auto-fluorescent property of pyoverdine and measured its production level through excitation at 400 nm and emission at 460 nm, in iron-limited medium. This assay revealed that 28 out of the 315 isolates produced no or less than 5% of the pyoverdine produced by characterized laboratory *Pseudomonas* reference strains (e.g. including *P. aeruginosa* PAO1, *P. protegens* CHA0, *P. putida* IsoF; see detailed description in Butaitė et al. 2017). We considered these 28 isolates as pyoverdine non-producers and thus potential cheaters. Furthermore, there was high variation in the pyoverdine levels among producers, with many isolates producing less than 50% of the pyoverdine produced by the reference strains. We considered this latter group of isolates as pyoverdine low-producers, which could be partial cheaters (Ghoul et al. 2014). We henceforth abbreviate non- and low-producers as NLPs.

For the current paper, we picked one or two NLPs (upon availability) from each community and used them as focal isolates to test whether they are locally adapted to exploit pyoverdine from producers. Overall, we had 14 non-producers and 10 low-producers, originating from seven soil communities (i.e. one soil community featured no NLPs) and eight pond communities. We paired each of the 24 focal isolates with four random producers from each of the four distance categories (i.e. same, close, intermediate, or distant community; 16 producers per focal strain). For producers to be included, they had to: (i) grow better than the corresponding NLPs under iron-limited conditions; (ii) produce more pyoverdine than the NLPs they were combined with; and (iii) differ in the *rpoD* sequence from the other three producers of the same community, and thus represent phylogenetically different strains. These criteria were met for all but one focal isolate that had to be paired with three producers with identical *rpoD* sequences form the same community, as no other options were available. This design resulted in 384 NLP-producer combinations (192 per habitat type). Given the limited number of producers available, some of them were combined with multiple focal isolates, so that our design finally featured 179 different pyoverdine producers (92 and 87 isolates from soil and pond, respectively).

### Supernatant assay

To quantify the extent to which NLPs can use pyoverdine from producers, we harvested pyoverdine-containing supernatants from all the producers and fed them to the focal NLPs. We followed the protocol described in our previous study (Butaitė et al. 2018). Specifically, we grew all producers (from overnight lysogeny broth cultures) in CAA medium (5 g casamino acids, 1.18 g K_2_HPO_4_·3H_2_O, 0.25 g MgSO_4_·7H_2_O per litre) supplemented with 200 μM 2,2’-dipyridyl as the iron chelator to stimulate pyoverdine production. Producers were grown in a total volume of 2 ml in 24-well plates, static at 25°C for 18 h. Subsequently, we centrifuged cultures for 10 min at 3,500 rpm (Eppendorf Centrifuge 5804R) and transferred 900 μl of the supernatants to AcroPrep Advance 96-well 1 ml filter plates (with a 0.2 μm supor membrane; Pall Corporation, USA), attached to an autoclaved 1.2 ml 96-well PCR plate (VWR). We centrifuged the samples in the filter plates together with the collection plates for 15 min at 2,500 rpm. The collection plates with sterile supernatants were sealed with Greiner SILVERseals and stored at −20°C.

Next, we grew the NLPs in 200 μl of LB in 96-well plates overnight static at 25°C. Overnight cultures of all focal strains were then adjusted to OD600 = 0.05 (optical density at 600 nm; measured with the microplate reader Infinite M200, Tecan Group Ltd., Switzerland). 2 μl of these adjusted cultures were transferred to four different variants of the CAA medium: (i) 180 μl CAA supplemented with 200 μM 2,2’-dipyridyl and 20 μl producer supernatant; (ii) 180 μl CAA supplemented with 200 μM 2,2’-dipyridyl and 20 μl CAA that underwent the same treatment as the supernatants including filtering and freezing; (iii) 180 μl CAA supplemented with 40 μM FeCl3 and 20 μl producer supernatant; (iv) 180 μl CAA supplemented with 40 μM FeCl3 and 20 μl CAA that underwent the same treatment as the supernatants including filtering and freezing. While (i) is our main treatment to quantify the effect of supernatants on focal strain growth, relative to the growth in non-supplemented medium (ii), treatments (iii) and (iv) serve as controls to assess the effect of supernatants on growth under iron-rich conditions where pyoverdine is not important for iron acquisition.

Each treatment was repeated four times for each strain combination. The plates were incubated statically for 15 h at 25°C. The final OD600 of cultures was measured using the microplate reader. We then used the OD600 measurements to calculate the “supernatant effect” under iron-limited conditions as [OD600 from treatment (i)] / [OD600 from treatment (ii)], and under iron-rich conditions as [OD600 from treatment (iii)] / [OD600 from treatment (iv)]. All ratios were log-transformed and values larger or smaller than zero indicate growth stimulation or inhibition, respectively. One focal strain did not grow under iron-limited conditions and the supernatant effect thus remained undefined. This strain had to be excluded from all subsequent analyses.

### Competition assays

Next, we directly competed NLPs against pyoverdine producers under iron-limited conditions to test whether the pyoverdine-mediated growth effects (as measured by the supernatant assay) translate into relative fitness consequences for the interacting strains. In order to distinguish the two competing strains, we integrated a single copy of a constitutively expressed mCherry marker into the chromosome of NLPs using the mini-Tn7 system (Choi and Schweizer 2006). We used both the electroporation and conjugation protocol by Choi & Schweizer (2006) with modifications described in Butaitė et al. (2017) for the successful tagging of 12 NLPs (six soil and six pond isolates), originating from five soil and five pond communities. Each of the 12 tagged isolates was competed against the corresponding 16 pyoverdine producers (four from each distance category) used for the supernatant assay. This resulted in 192 NLP-producer combinations (96 per habitat type).

The competition experiments entailed the following steps: (i) We grew isolates as monocultures overnight in 200 μl LB medium in 96-well plates for about 17 h at 25°C. (ii) We adjusted the cultures to OD600 = 0.05 in LB. (iii) We mixed strains in a 1:5 volumetric ratio (20 μl of mCherry-tagged NLPs culture with 100 μl of the untagged producer culture). (iv) We transferred 2 μl of strain mixes, but also monocultures and media blanks as controls, to 200 μl CAA containing 200 μM 2,2-dipyridyl in 96-well plates, in five-fold replication. (v) We let strains compete for about 46 h at 25°C under static conditions. (vi) We used flow cytometry to estimate the initial and final frequencies of the competing strains (see detailed method below). Important to note is that we used filtered medium (passed through a 0.22 μm filter) for all competition experiments to reduce the medium background signal during flow cytometry measurements. (vii) We estimated the relative fitness of the NLPs as: *v* = [a_t_(1 − a_0_)]/[a_0_(1 − a_t_)], where a_0_ and a_t_ are initial and the final frequencies of NLPs in the mixed cultures with the producer, respectively (Ross-Gillespie et al. 2007). We log-transformed *v* values, whereby *v* < 0 indicates a decrease and *v* > 0 an increase in the relative fitness of NLPs compared to its pyoverdine-producing competitor.

### Flow cytometry

Prior to flow cytometry measurements, we fixed cells (to stop any biological process) using paraformaldehyde. We first dissolved paraformaldehyde in PBS (phosphate-buffered saline) to obtain a 10% stock solution, and then further diluted the stock with 0.85% NaCl to obtain a final concentration of 2%. This solution was filter-sterilized using a 0.22 μm filter. We used the paraformaldehyde-PBS-NaCl solution to fix and dilute both mono- and mixed cultures before (a representative aliquot) and after the competition period. Appropriate dilution factors were chosen to obtain less than 10,000 events/s, which is the upper limit our flow cytometer (LSR II Fortessa, BD Biosciences) could handle. Cells were fixed for 20 min in the dark at room temperature, and then kept at 4°C prior to staining. We stained fixed cells with Sybr Green I (Invitrogen; the commercial stock was diluted by 2*10^4^x) for 30 min at room temperature in the dark. Pre-competitions (mono- and mixed cultures) were analysed in triplicates, while competitions involved five independent replicates per strain combination. We used the HTS mode (automated 96-well sampler) of the flow cytometer to assess strain frequencies and total cell counts in 10 μl per sample, at the flow cytometry core facility of the University of Zurich. Sybr Green I (excitation: 488 nm / emission: 530 nm) and mCherry (excitation: 561 nm / emission: 610/20 nm) fluorescence was measured using the corresponding detection filters. We used the FlowJo software (Tree Star) for data gating and the calculations of strain frequencies. Because the mCherry signal was typically too weak in the pre-competition samples, we used Sybr Green I to assess cell counts in monocultures prior to mixing and used those to infer strain frequencies in the 1:5 volumetric mixes used to initiate the competitions. After the competition period, the mCherry signal was strong, which allowed an unambiguous distinction between tagged and untagged cells.

### Statistical analysis

We first explored whether phylogenetic relatedness varied with geographical distance in our full collection of soil and pond isolates. To account for the possibility that phylogenetic diversity changes more distinctively over spatial scales in the more structured soil environment as compared to the more diffusive pond environment, we performed separate Mantel tests for soil and pond isolates. To further assess whether relatedness between NLP and producer pairs used in our experiments changes with distance, we fitted a linear mixed model (LMM) using phylogenetic relatedness as response variable, and habitat (soil or pond), distance category (same, close, intermediate or far), and their interaction as explanatory variables. To explore the potential determinants of the supernatant effect, we fitted a second LMM using the log-transformed supernatant effect as continuous response variable, and habitat, distance category, iron-availability (iron-limited or iron-replete), phylogenetic relatedness (centered and scaled to unit variance) as well as all their interactions as explanatory variables. To finally assess the determinants of the outcome of direct competition between NLPs and producers, we fitted a third LMM using the (log-transformed) relative fitness values of the NLP as continuous response variable, and habitat, distance, relatedness, and their interactions as explanatory variables. Additionally, this third LMM contained the (log-transformed) supernatant effect and the initial ratio of the NLP as main effects to assess whether the supernatant effect was predictive of competitive outcomes, and whether these outcomes depended on variation in the relative starting frequency of the NLPs, respectively.

In our experiments NLPs interacted with multiple pyoverdine producers and some pyoverdine producers were paired with multiple NLPs. To account for the resulting non-independent repeated measurements, we initially fitted all models as random intercept models using NLP identity nested within their community, and producer identity nested within producer community as random effects. We then simplified models in a two-step procedure. First, we simplified the random component of the model based on likelihood ratio tests of model reduction. In a second step, we simplified the fixed component of each model by dropping non-significant interaction terms (p > 0.05). All statistical analyses were conducted using the statistics software R version 3.5.0 (www.r-project.org). Mantel tests were performed using the ‘mantel’ function of the *ecodist* package with 9999 permutations and 999 iterations for the bootstrapped confidence limits. Mixed models were implemented using the ‘lmer’ function of the *lme4* package. The p-values of effects in these models were obtained using the ‘Anova’ function of the *car* package and the ‘summary’ function of the *lmerTest* package.

## RESULTS

### *Pseudomonas* genetic relatedness varies across space in soil, but not in pond

In a first analysis, we compared the phylogenetic relatedness between pairs of isolates based on *rpoD* sequence similarities across the four geographical distance categories of our sampling scheme (Fig. 1A). When including all the 304 isolates with a sequence length > 500 bp, we found that phylogenetic relatedness decreased with geographical distance among soil isolates (Mantel R [upper CI; lower CI] = -0.076 [-0.108; -0.052], p = 0.0001; Fig. 1B), but was independent of distance among pond isolates (Mantel R [upper CI; lower CI] = -0.008 [-0.017; -0.001], p = 0.2019; Fig. 1B). When restricting the analysis to the isolate pairs we used for the supernatant assay, the phylogenetic relatedness did not differ between habitats and was overall independent of distance (LMM: habitat: χ^2^_1_ = 1.472, p = 0.225; distance: χ^2^_3_ = 2.874, p = 0.412; Fig. 1C), a result that can likely be attributed to the reduced sample size and thus lower statistical power. Overall, our results show that there is weak but significant phylogenetic structuring of the *Pseudomonas* communities across a distance of 50 m in the soil, but not in the pond habitat.

### Phylogenetic relatedness is the main predictor of the supernatant effect, yet geographical distance also plays a role among pond isolates

To test whether pyoverdine-meditated social interactions vary across geographical distance, we fed pyoverdine-containing supernatants from 175 different producers to 23 focal pyoverdine non- and low-producers (NLPs), such that each NLP received supernatants from 16 producers originating from the four distance categories (Fig. 1A; four producers per each distance category). Under iron-limited conditions where pyoverdine is important for growth, we observed 122 cases (soil: 56, pond: 66) in which the supernatant of producers stimulated the growth of NLPs (Fig. 2). Because we know from our previous work that supernatant effects are predominantly driven by pyoverdine under iron limitation (Butaitė et al. 2017), our results indicate that many NLPs can take up heterologous pyoverdines to overcome iron limitation. Conversely, we found 246 cases (soil: 120, pond: 126) in which the supernatant of producers inhibited the growth of NLPs (Fig. 2), suggesting that these isolates lack the ability to use the specific heterologous pyoverdine types fed.

**Figure 2.**
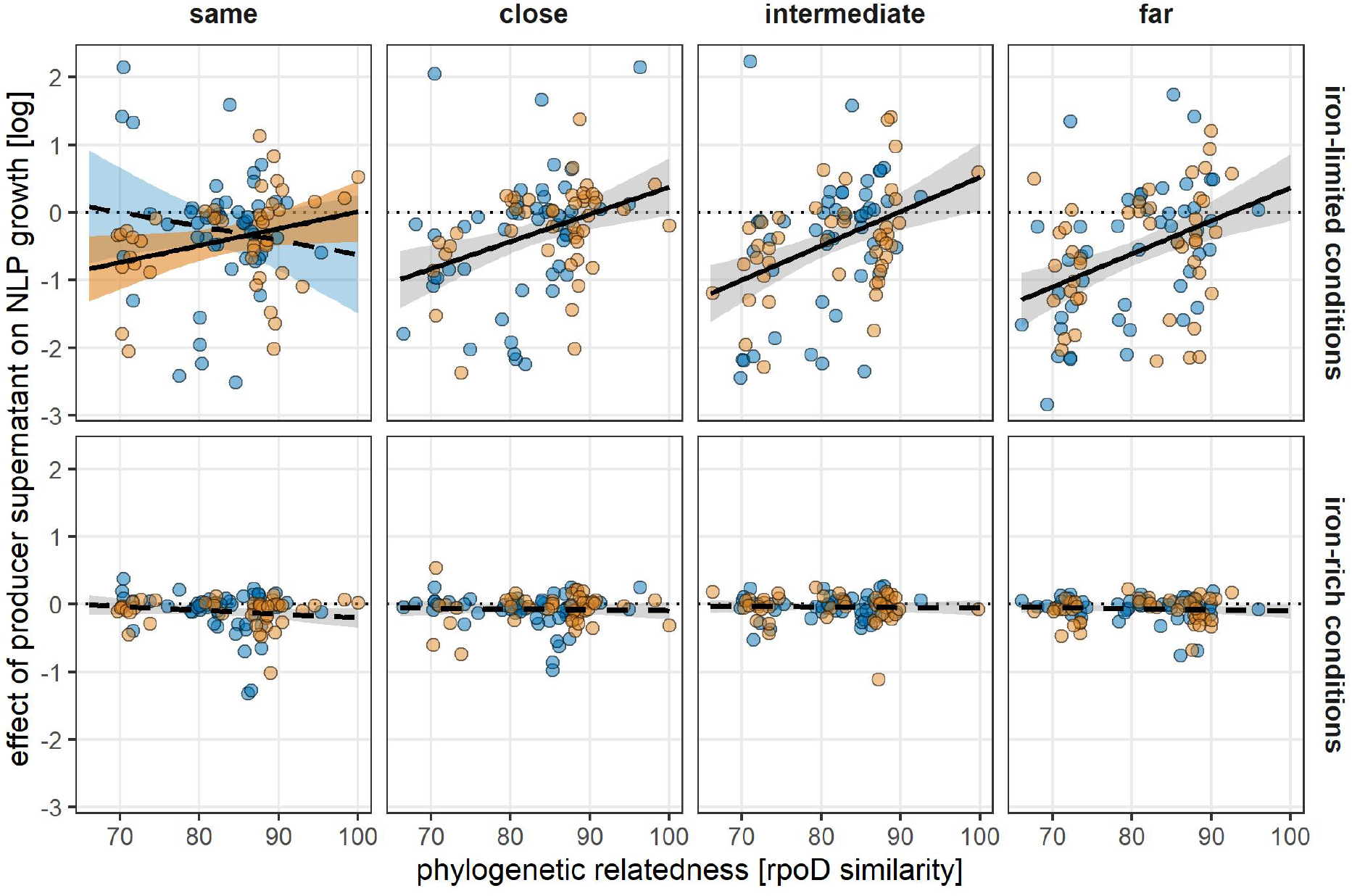
The supernatant effect correlates positively with the phylogenetic relatedness between interacting isolates across all geographical distance categories, except for interactions among pond isolates from the same community. Depicted is the relationship between phylogenetic relatedness (based on *rpoD* sequence similarities) between pairs of pyoverdine producers and NLPs and the supernatant effect under iron-limited and iron-rich conditions for soil (yellow) and pond (blue) isolates across the four different geographical distance categories. The solid and dashed lines indicate significant and non-significant relationships, respectively. Dotted horizontal lines indicate the null line where supernatants have no effect on the growth of NLPs. Shaded areas are 95% confidence intervals; grey = relationship applying to both soil and pond isolates, yellow = soil-specific relationship, blue = pond-specific relationship.

When comparing these results across treatments and conditions, we found that the supernatant effect was shaped by an interaction between habitat, geographical distance, and the relatedness between producer and NLP (Table S1). We therefore split the model and analyzed the supernatant effect separately for soil and pond isolates. Among soil isolates, the supernatant effect was shaped by an interaction between iron limitation and phylogenetic relatedness (Table 1). Specifically, the supernatant effect increased with relatedness when iron was limited (slope ± SE: 0.274 ± 0.036, t_337.1_ = 7.584, p < 0.001; Fig. 2), but was independent of relatedness under iron-replete conditions (0.022 ± 0.036, t_377.1_ = 0.619, p = 0.536; Fig. 2). By contrast, the supernatant effect did not vary with geographical distance (Table 1, Fig. 2). These findings suggest that there is no local adaptation with regard to pyoverdine-mediated social interactions (ranging from growth stimulation to inhibition) in *Pseudomonas* soil communities, but that interaction patterns are driven by phylogenetic relatedness between the isolates.

**Table 1.**
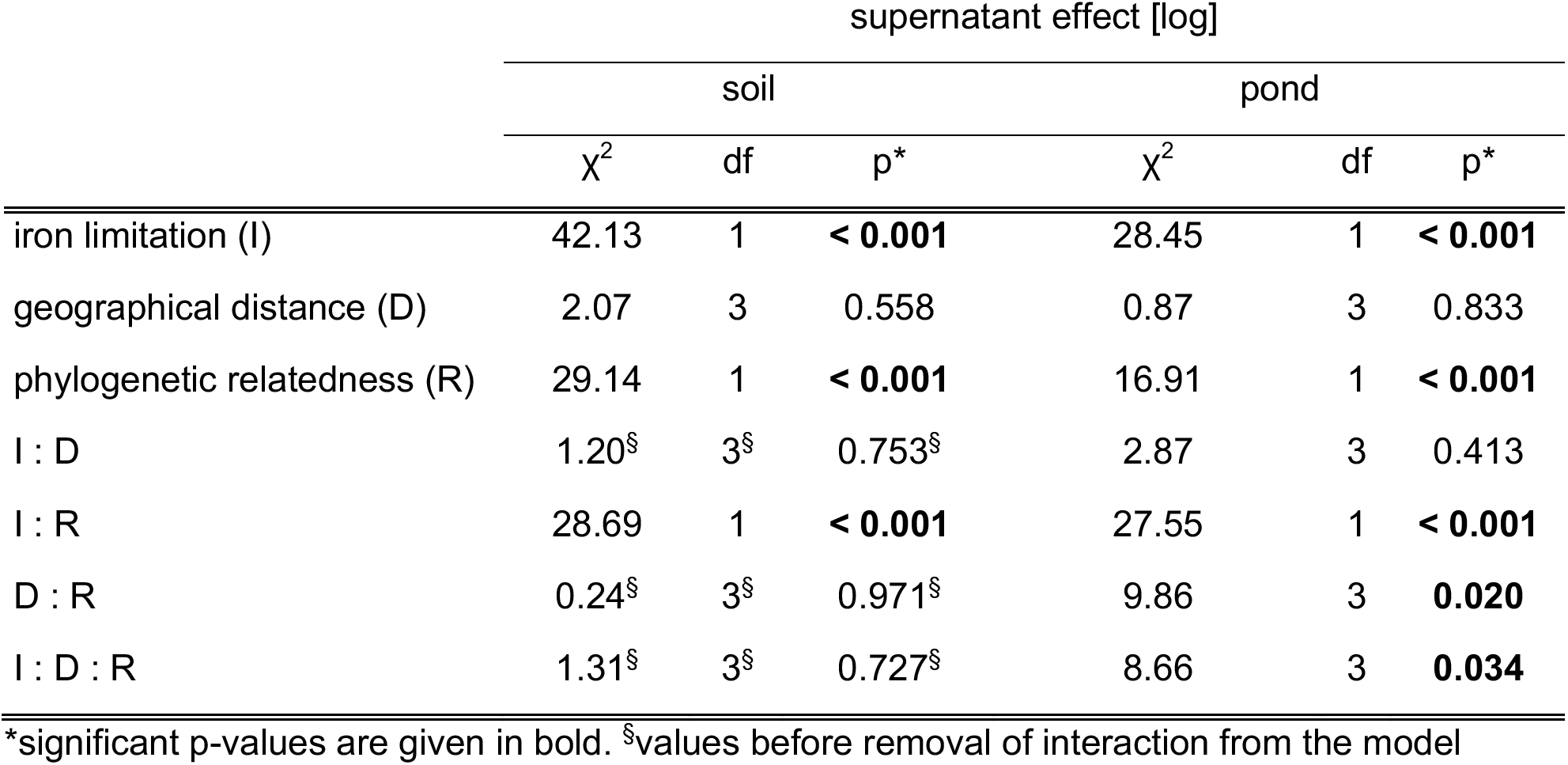
Statistical analysis of the supernatant effect on NLPs. Influence of habitat (soil vs. pond), iron limitation, geographical distance, and phylogenetic relatedness on the (log-transformed) effect of the supernatant of pyoverdine producers on the growth of pyoverdine non- and low-producers (NLPs).

|  | supernatant effect [log] |  |  |  |  |  |
| --- | --- | --- | --- | --- | --- | --- |
|  | soil |  |  | pond |  |  |
| | $\chi^2$ | df | p* | $\chi^2$ | df | p* |
| iron limitation (I) | 42.13 | 1 | **< 0.001** | 28.45 | 1 | **< 0.001** |
| geographical distance (D) | 2.07 | 3 | 0.558 | 0.87 | 3 | 0.833 |
| phylogenetic relatedness (R) | 29.14 | 1 | **< 0.001** | 16.91 | 1 | **< 0.001** |
| I : D | 1.20<sup>§</sup> | 3<sup>§</sup> | 0.753<sup>§</sup> | 2.87 | 3 | 0.413 |
| I : R | 28.69 | 1 | **< 0.001** | 27.55 | 1 | **< 0.001** |
| D : R | 0.24<sup>§</sup> | 3<sup>§</sup> | 0.971<sup>§</sup> | 9.86 | 3 | **0.020** |
| I : D : R | 1.31<sup>§</sup> | 3<sup>§</sup> | 0.727<sup>§</sup> | 8.66 | 3 | **0.034** |

\*significant p-values are given in bold. <sup>§</sup>values before removal of interaction from the model

The patterns among pond isolates differed from those among soil isolates in one notable aspect (Fig. 2): the supernatant effect depended on an interaction between iron limitation, phylogenetic relatedness, and geographical distance (Table 1). This interaction arose because under iron-limited conditions, the supernatant effect was independent of phylogenetic relatedness among isolates from the same pond community (-0.056 ± 0.158, t_187.4_ = -0.354, p = 0.724), but increased with phylogenetic relatedness among isolates from close (0.469 ± 0.125, t_175.9_ = 3.761, p < 0.001), intermediate (0.492 ± 0.146, t_184.1_ = 3.359, p < 0.001), and far communities (0.509 ± 0.123, t_188.3_ =4.154, p < 0.001; Fig. 2). These results show that the otherwise predominant effect of phylogenetic relatedness on the supernatant effect is eroded among members of the same pond community. Similar to the soil isolates, the supernatant effect was neither linked to phylogenetic relatedness (χ^2^ = 0.15, df = 1, p = 0.696) nor geographical distance (χ^2^ = 6.80, df = 3, p = 0.075) under iron-replete conditions.

### The relative fitness of pyoverdine NLPs in direct competition with producers correlates with the results of the supernatant assay

To examine whether the pyoverdine-mediated growth effects in the supernatant assay translate into relative fitness consequences in direct competitions between pyoverdine NLPs and producers, we competed the two types of isolates against each other using the same spatially arranged design, albeit with a reduced sample size. Overall, we had 192 producer-NLP combinations (soil: 96, pond: 96), where each of the 12 NLPs was competed against 16 producers originating from the four distance categories (Fig. 1A; four producers per each distance category).

We observed that competitions between pyoverdine producers and NLPs were mostly won by producers. In only 23 cases we observed a relative fitness increase of the NLPs, whereas producers won in 169 cases. Overall, the relative fitness patterns were similar to those observed for the supernatant effects (Fig. 3A). The relative fitness of NLPs increased with phylogenetic relatedness (0.263 ± 0.117, t_155.0_ = 2.255, p = 0.024), but neither varied with geographical distance nor between habitats (Table 2). Moreover, the relative fitness of the NLPs correlated positively with the supernatant effect (0.792 ± 0.104, t_165.1_ = 7.636, p < 0.001; Fig. 3B). These findings show that phylogenetic relatedness is the main factor determining the outcome of competition for iron between pseudomonads within and across communities. Moreover, they show that the pyoverdine-mediated growth effect in the supernatant assay translates into relative fitness consequences, which highlights the key role of siderophores in determining the outcome of strain interactions.

**Table 2.**
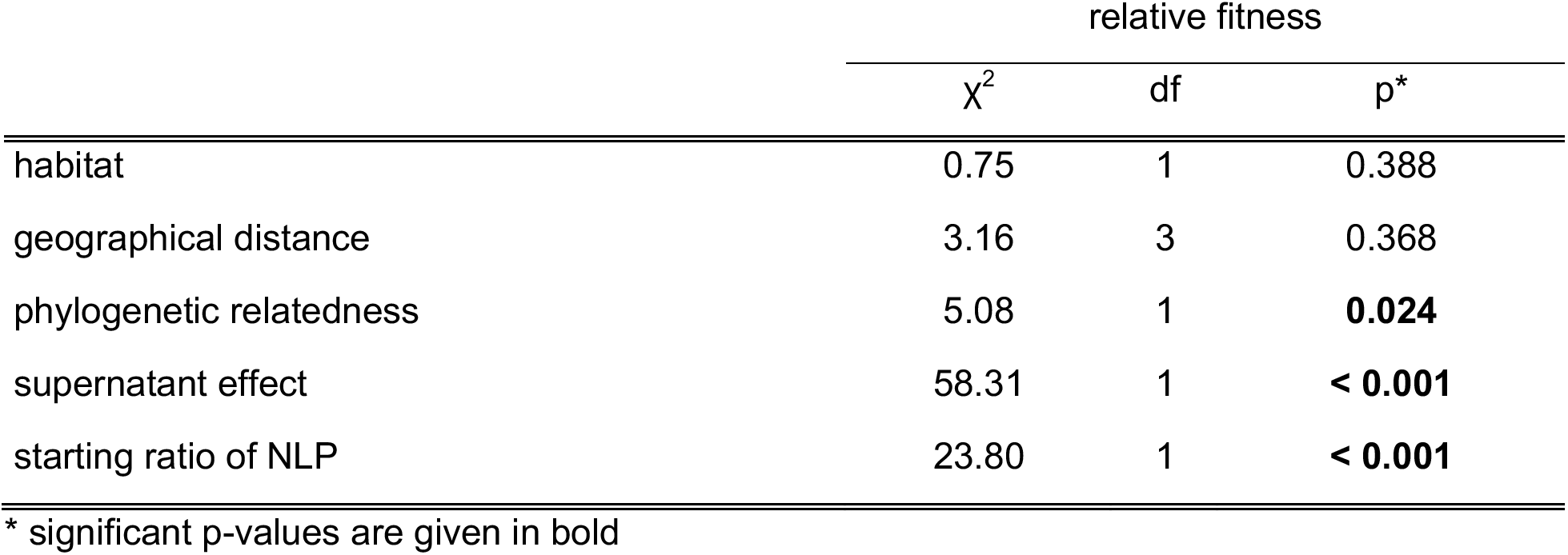
Statistical analysis of the relative fitness of NLPs in competition with pyoverdine producers. Influence of habitat (soil vs. pond), geographical distance, phylogenetic relatedness, the (log-transformed) supernatant effect (as shown in Fig. 2), and the starting ratio of the NLPs on their relative fitness in competition with producers.

**Figure 3.**
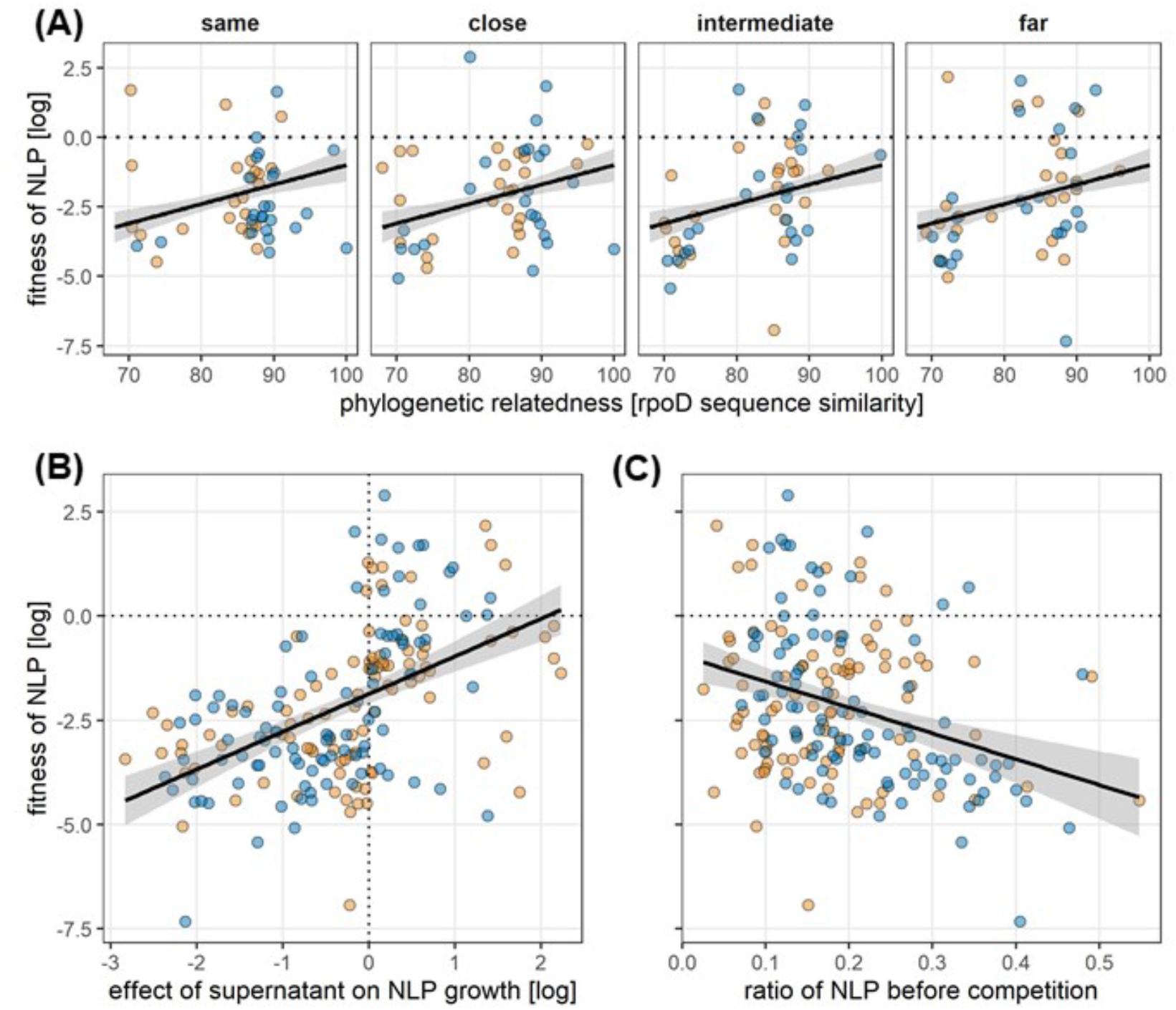
Competitive outcomes between pyoverdine producers and NLPs depend on phylogenetic relatedness, but do not differ across the four geographical distance categories. (A) The relative fitness of pyoverdine non- and low-producers (NLPs) increases with the phylogenetic relatedness between producer and NLP but does not differ between the four different distance categories in soil (yellow) and pond (blue). (B) The relative fitness of NLPs correlates positively with the supernatant effect. (C) The relative fitness of NLPs decreases the more common (i.e. higher in frequency) they were at the beginning of the competition. Solid lines depict significant relationships. Shaded areas are 95% confidence intervals. Dotted horizontal lines indicate equal fitness of producers and NLPs. The dotted vertical line in (B) indicates no effect of the supernatant on the growth of NLPs.

Finally, the relative fitness of NLPs was also dependent on their initial frequency in the population (Fig. 3C). In particular, we found that the relative fitness of NLPs increased the rarer they initially were (-5.930 ± 1.216, t_133.1_ = -4.878, p < 0.001; Fig. 3C), a pattern predicted by social evolution theory for microbes (Ross-Gillespie et al. 2007). It arises because non-producers can exploit public goods more efficiently when surrounded by many producers that deliver the exploitable good. We could test for this pattern because, although we adjusted all strains to OD600 = 0.05 prior to the volumetric mixing of 1 NLP: 5 producer units, the actual strain frequency (assessed with flow cytometry) of NLPs varied considerably: 0.025-0.55.

## DISCUSSION

We set out to test whether siderophore-mediated cooperation and cheating can spur patterns of local adaptation in soil and pond communities of *Pseudomonas* bacteria, sampled across four geographical distance categories. We argued that local adaptation could manifest because pyoverdine, the main siderophore of fluorescent pseudomonads, shows high inter-strain variability in its molecule structure and in the receptor required for siderophore uptake (Ghysels et al. 2004; Smith et al. 2005; Meyer et al. 2008; Butaitė et al. 2017). Consequently, we hypothesized that pyoverdine non- and low-producers (NLPs) might become adapted to efficiently exploit pyoverdines produced by local producers and/or producers might evolve strategies to particularly resist cheating by local non-producers. While we indeed observed that the level of pyoverdine exploitation and cheating resistance varied considerably between interacting strain pairs, we found only little evidence that this variation correlated with geographical distance. Instead, there was a strong signature of phylogenetic relatedness, whereby NLPs were generally better at exploiting the pyoverdines of more closely related producers, irrespective of whether strain pairs originated from the same, close or more distantly related communities. There was one notable exception: within local pond communities the relationship between phylogenetic relatedness and pyoverdine-mediated growth effects was broken. In the sections below, we first discuss possible reasons for the overall weak evidence of local adaptation and then turn to the special interaction pattern observed in local pond communities. Finally, we discuss reasons for why phylogenetic relatedness could be such an important factor driving pyoverdine-mediated social interactions.

There are a number of ecological and evolutionary forces that are known to prevent local adaptation (Kawecki and Ebert 2004; Blanquart et al. 2013; Savolainen et al. 2013). First, high dispersal rates of individuals across the geographical scale sampled could promote gene flow between patches and erode any form of local adaptation. This scenario might apply to the pond habitat, where we found no significant genetic structuring across the geographical scale sampled (Fig. 1B). Conversely, significant genetic structuring occurred in soil, suggesting that gene flow is limited in this habitat. Second, local adaptation cannot occur if there is no genetic variation at the trait of interest. This scenario does certainly not apply in our case because our own data show tremendous variation in pyoverdine-mediated growth inhibition and stimulation (Fig. 2). Moreover, our previous genetic analysis on the pyoverdine system of a small subset of pond and soil isolates revealed high pyoverdine molecule and receptor diversity, thus providing enough variability for natural selection to act on (Butaitė et al. 2017). Third, local adaptation requires natural selection to be the driving force of evolution. When other factors, such as genetic drift, play a major role then local adaption cannot occur. While we typically assume that bacterial population sizes are high enough for natural selection to operate, we know too little on the effective size of our *Pseudomonas* populations to firmly exclude genetic drift as a factor preventing local adaptation. Fourth, frequent temporal variation in environmental conditions can undermine local adaptation. The logic is simple: a beneficial strategy in response to condition A might become unfavorable when conditions change to B. It is easy to see how this scenario can apply to fluctuations in abiotic environmental factors. For example, fluctuation in iron availability might affect the selection pressure on NLPs. Under iron-rich conditions, NLPs are not dependent on pyoverdine producers, such that any type of NLPs would be selectively favored, which would undermine local adaptation. In contrast, NLPs depend on producers and a suitable pyoverdine receptor array under iron-limited conditions, which could favor local adaptation. Our previous work revealed that iron availability was consistently low in pond communities, but much higher yet more variable in soil communities (Butaitė et al. 2018). Taken together, we reason that increased gene flow in pond and higher environmental fluctuations in soil could be likely explanations for the absence of local adaptation patterns.

We now turn to the local pond communities, where the otherwise prevalent positive relationship between the supernatant effect and the phylogenetic relatedness was absent (Fig. 2). This finding recovers our previous result on within-community interaction patterns (Butaitė et al. 2017). In this previous paper, we further investigated the genetic basis of pyoverdine-mediated interactions by sequencing the genomes of 24 *Pseudomonas* isolates. One of the key findings was that most isolates had more than one gene encoding for a pyoverdine receptor (median receptor homologues per isolate = 4). We speculated that (i) horizontal gene transfer could enable isolates to acquire receptor variants of pyoverdine types they do not produce themselves, and (ii) frequent horizontal gene transfer would erode the signature of phylogenetic relatedness. We then proposed that horizontal gene transfer is likely to happen more often in the diffusive pond environment, which can explain the absence of an effect of phylogenetic relatedness at the local population scale in pond (Butaitė et al. 2017). In further support of horizontal gene transfer being more prominent in the pond habitat, we found that pond isolates had significantly higher numbers of pyoverdine receptor homologues than soil isolates (median values 5.5 vs. 2.5; Wilcoxon rank sum test: W = 33, p = 0.0243). These observations are interesting, but do they support a scenario of local adaptation? They could indeed do so, especially in the case where horizontal gene transfer predominantly occurs within a community (Cordero and Polz 2014). Then, any NLP could quickly acquire the receptors of the prevalent producers via horizontal gene transfer. But why were the NLPs then not better at exploiting the local versus the more distant producers? One plausible answer is that the rate of adaptation is similar between NLPs and pyoverdine producers, so that there is simply no overall winner in the antagonistic arms race (Kümmerli et al. 2015), and traces of local adaptation remain thus masked (Kaltz and Shykoff 1998).

An important question to address by any study on local adaptation is whether we have looked at the biologically relevant spatial scale. For a *Pseudomonas* bacterium with a length between 1 to 5 μm, a geographical scale of 50 m seems large. For the soil habitat at least, it indeed seems that gene flow is restricted across this scale. However, we also need to examine the lower end of the scale and ask whether isolates from the ‘same’ community would indeed interact in nature. We do not have a conclusive answer to this question. All we know is that pseudomonads are typically motile (Sampedro et al. 2015), and that pyoverdine secreted on surfaces can be shared between single cells across a range of 100 μm (Weigert and Kümmerli 2017), a range that can be much larger between macroscopic colonies (Kümmerli et al. 2009a). It thus seems reasonable to assume that at least a fraction of strains isolated from our 2 cm^3^ soil cores might be able to interact with each other in nature. When comparing to other studies, it becomes evident that the scale across which geographical effects are observed varies between bacterial species and habitats, and can cover any range from the centimeter to the kilometer scale (Vogel et al. 2003; Vos and Velicer 2008; Vos and Velicer 2009; Hawlena et al. 2010; Hawlena et al. 2012; Kraemer et al. 2016; Bruce et al. 2017b; Kraemer et al. 2017).

A key finding of our study is that the phylogenetic relatedness between pyoverdine NLPs and pyoverdine producers is the main predictor of (i) the extent to which NLPs can benefit from the pyoverdine secreted by producers (Fig. 2), and (ii) the relative fitness of the NLPs in direct competition with producers (Fig. 3A). Moreover, we observed that the effects of (i) and (ii) correlated positively with each other (Fig. 3B). The strong phylogenetic signature is perhaps surprising at first sight given that isolates have many different pyoverdine receptors (Butaitė et al. 2017), which should allow them to use a range of different pyoverdine types and not only those from close relatives (Sexton et al. 2017). However, our results now indicate that both absolute and relative fitness consequences seem to be largely driven by the cognate pyoverdine receptor. In other words, NLPs lost or reduced the ability to produce pyoverdine, but kept their cognate receptor (Butaitė et al. 2017), and they seem to be most efficient in using the pyoverdine they once produced themselves and is still produced by close relatives. The additional pyoverdine receptors, possibly acquired through horizontal gene transfer, might also be beneficial, but their contribution to fitness seems to be smaller and do not override the effect of phylogenetic relatedness in most cases, except within local pond communities.

Although our main results are based on supernatant assays, involving pyoverdine but also other secreted compounds, we are confident that the measured fitness effects are indeed predominantly due to pyoverdine. Our reasoning is based on the fact that all strong supernatant effects (inhibition, stimulation, positive correlation with phylogenetic relatedness) disappeared when the experiment was repeated in iron-rich medium (Fig. 2). Under these conditions, pyoverdine is no longer produced (Tiburzi et al. 2008; Kümmerli et al. 2009b), while other secreted compounds potentially influencing fitness still are. Moreover, we have previously purified pyoverdine and demonstrated that the supernatant effect strongly correlates with the fitness effects of the actual pyoverdine (Butaitė et al. 2017).

In summary, our study contributes to the increasing body of evidence that siderophores are important compounds driving species interactions in a variety of habitats, and in determining the composition and assembly of natural bacterial communities (Cordero et al. 2012; Bruce et al. 2017a; Butaitė et al. 2017; Gu et al. 2020; Kramer et al. 2020b). While local adaptation might play a minor role in defining interaction patterns, we show that pyoverdines have strong yet variable effects on bacterial fitness: (i) the effect of heterologous pyoverdines on NLP fitness ranges from growth inhibition to promotion; (ii) the increased ability to use pyoverdine from other *Pseudomonas* isolates translates into higher relative fitness in direct competition with producers; and (iii) the relative fitness of NLPs was highest when they were rare, demonstrating that negative frequency-dependent fitness patterns occur in natural communities.

## Data availability statement

All the raw data of this study will be made available on the Dryad depository upon the acceptance of this article for publication.

## Acknowledgements

We thank the flow cytometry facility of the University of Zurich and Peter J. Richards for technical support, and Stefan Wyder, Sébastien Wielgoss, Michael Baumgartner and Marta Pinto Carbó for bioinformatical support. This work was funded by the Swiss National Science Foundation (grants no. 165835 and 182499 to RK), the European Research Council (under the European Union’s Horizon 2020 programme, grant agreement no. 681295 to RK), the German Science Foundation (DFG; KR 5017/2-1 to JK), and the Forschungskredit of the University of Zurich (FK-15-082 to EB).

## Author contributions

EB and RK designed the experiments; EB conducted the experiments; EB and JK analyzed the data; EB, JK and RK interpreted the data and wrote the paper.

## Supplementary material

**Table S1.** Supernatant effect (full model). Influence of habitat, iron limitation, geographical distance, and phylogenetic relatedness on the (log-transformed) effect of the supernatant on the growth of pyoverdine non-/low-producers.

|  | supernatant effect [log] |  |  |
| --- | --- | --- | --- |
| | $\chi^2$ | df | p |
| habitat | 0.20 | 1 | 0.659 |
| iron limitation | 68.87 | 1 | **< 0.001** |
| geographical distance | 2.35 | 3 | 0.503 |
| phylogenetic relatedness | 39.00 | 1 | **< 0.001** |
| habitat : iron limitation | 0.75 | 1 | 0.388 |
| habitat : geographical distance | 0.24 | 3 | 0.971 |
| iron limitation : geographical distance | 3.77 | 3 | 0.288 |
| habitat : phylogenetic relatedness | 0.39 | 1 | 0.535 |
| iron limitation : phylogenetic relatedness | 52.55 | 1 | **< 0.001** |
| geographical distance : phylogenetic relatedness | 4.35 | 3 | 0.227 |
| habitat : geographical distance : phylogenetic relatedness | 8.03 | 3 | **0.045** |

\* significant p-values are given in bold print

